# Poxviruses package viral redox proteins in lateral bodies and modulate the host oxidative response

**DOI:** 10.1101/2020.12.09.418319

**Authors:** Susanna R. Bidgood, Karel Novy, Abigail Collopy, David Albrecht, Melanie Krause, Jemima J. Burden, Bernd Wollscheid, Jason Mercer

**Affiliations:** MRC Laboratory for Molecular Cell Biology, University College London, Gower Street, London, WC1E 6BT; Institute of Molecular Systems Biology, ETH Zürich / HPT E52.1, Auguste-Piccard-Hof 1, CH-8093 Zürich, Switzerland; Swiss Institute of Bioinformatics (SIB), Switzerland; Institute of Microbiology and Infection, University of Birmingham, Birmingham, United Kingdom

**Keywords:** Vaccinia, poxvirus, lateral body, ROS, spatial proteotyping

## Abstract

All poxviruses contain a set of proteinaceous structures termed lateral bodies (LB) that deliver viral effector proteins into the host cytosol during virus entry. To date, the spatial proteotype of LBs remains unknown. Using the prototypic poxvirus, vaccinia virus (VACV), we employed a quantitative comparative mass spectrometry strategy to determine the poxvirus LB proteome. We identified a large population of cellular proteins, the majority being mitochondrial, and 15 viral LB proteins. Strikingly, one-third of these comprise the full set of VACV redox proteins whose LB residency could be confirmed using super-resolution microscopy. We further show that VACV infection exerts an anti-oxidative effect on host cells and that artificial induction of oxidative stress impacts early gene expression and virion production. In addition to defining the spatial proteotype of these enigmatic viral structures, these findings implicate poxvirus redox proteins as modulators of host oxidative anti-viral responses and provide a solid starting point for future investigations into the role of LB resident proteins in host immunomodulation.

## Introduction

The main goal of all viruses is to transfer a replication competent genome from one host cell to another (1, 2). This involves the delivery of the viral genome and accessory proteins to the cytosol or nucleus. In doing so they trigger cellular innate immune responses, a vast network of receptors and signalling cascades designed to prevent pathogen infection and replication (3, 4). In turn viruses have evolved numerous mechanisms to evade and manipulate these host immune defences (5, 6). While the majority of viruses appear to do this through the expression of nascent proteins during replication, both alphaherpesviruses and poxviruses have been reported to package and subsequently deliver accessory proteins with immunomodulatory capacity (7–11).

Herpesviruses do this using tegument, a protein layer between the capsid and membrane, which slowly dissociates during transit of viral capsids through the cytoplasm *en route* to the nucleus (9, 12–15). Poxviruses achieve this through two proteinaceous viral substructures termed lateral bodies (16–18). Structurally, pox virions are brick-shaped particles composed of three main substructures: the viral core which houses the dsDNA genome, the lateral bodies (LBs) which reside on either side of the core, and the viral membrane that encapsulates these structures (10, 19).

Vaccinia virus (VACV), the prototypic poxvirus, enters host cells by macropinocytosis (20–22). Upon fusion of viral and macropinosome membranes the viral core undergoes primary uncoating characterized by core expansion and the initiation of early gene transcription (16, 23). The LBs, which detach from the cores upon fusion, are deposited into the cytoplasm where they take on an effector function (24).

To date, attempts to define the complete composition of LBs and the function of their various protein constituents have been confounded by the molecular complexity of poxviruses (composed of more than 80 different viral proteins), and the inability to isolate intact LBs from viral particles or infected cells (8, 18, 25). Recently, we identified the first 3 *bona fide* poxvirus LB proteins: An abundant viral phosphoprotein F17R, the viral phosphatase H1, and a viral glutaredoxin-2, G4L (8). During late infection F17R acts to dysregulate (mTOR) and suppress interferon-stimulated gene responses (26, 27). As F17R accounts for 69% of LB mass, and its partial degradation mediates the release of the viral phosphatase H1L (8), we presume that F17R serves as the LB structural protein although any early host-modulatory role has yet to be investigated. The H1L phosphatase inhibits interferon-γ driven, anti-viral immunity through dephosphorylation of the transcription factor STAT1 (28, 29). This activity of H1L is dependent on proteasome-mediated disassembly of LBs and independent of viral early gene expression (8). G4L is a functional glutaredoxin whose activity is required for the formation of viral disulphide bonds during virion assembly (30–32). G4L is expressed late, packaged into LBs but has not been assigned any host-modulatory activity (8, 31).

Nevertheless, the generation of cellular reactive oxygen and nitrogen species (RONS) is an important component of cellular defense against incoming pathogens (33). All viral infections cause a redox imbalance in the host cell, the outcome of which depends on the duration and magnitude of this imbalance (34, 35). Most commonly a pro-oxidative effect is triggered, which initiates anti-viral responses including autophagy, cell death, mTOR inhibition, RONS-coupled pattern recognition receptor signalling (36). While some viruses (vesicular stomatis virus) (37, 38) are restricted by this, several have evolved ways to harness the oxidative environment (influenza, respiratory syncytial virus, and human immunodeficiency virus) (39–45), and others to combat it (human cytomegalovirus) (46, 47).

As a family of viruses that replicate exclusively in the cytoplasm, poxvirus replication is not surprisingly sensitive to RONS. IFNγ-driven nitric oxide (NO) production was shown to inhibit VACV and ectromelia virus replication in mouse macrophages (48), and NOS2-deficient mice are highly susceptible to ectromelia despite no known loss of ectromelia-targeted immune responses (49). These reports suggest that poxviruses must overcome RONS for successful replication. Interestingly, VACV encodes a set of redox proteins including a human homolog of glutaredoxin-1 (O2L), the previously mentioned glutaredoxin G4L which is a homolog of human glutaredoxin-2, a SOD-like dismutase (A45R), a thiol oxidoreductase (A2.5L), and a protein with 2 putative thioreductase CXXC motifs (A19L) (31, 50–53). While no immunomodulatory role has been elucidated, VACV mutants deleted for these various proteins indicate that A2.5L and G4L are required for cytoplasmic disulphide bond formation and virion assembly (30, 32, 52), and A19L for virus transcription and assembly (53, 54). Deletion of either A45R or O2L suggests that they are dispensable for VACV replication (51, 55).

Using controlled degradation of virions in combination with quantitative mass spectrometry-based proteotyping stratagies we define the viral and cellular proteins that constitute the VACV LB proteome. In addition to numerous cellular proteins, 15 viral LB proteins, including all VACV-encoded redox proteins were identified. Follow-up analysis of the impact of ROS on VACV infection indicated that VACV exerts an anti-oxidative effect on host cells and that high ROS levels block viral replication. In addition to elucidating the protein composition of poxvirus LBs, these findings further implicate LBs as early immunomodulatory delivery packets and provide a solid starting point for additional studies regarding the role of poxvirus redox proteins in modulating host oxidative responses to infection.

## Results

### Defining the poxvirus lateral body spatial proteotype

With a few exceptions, the compendium of proteins that make up poxvirus LBs remain unknown (7, 8). To define and investigate the LB proteome we employed sub-viral fractionation of VACV MVs in combination with quantitative mass spectrometry-based analysis (illustrated in Fig 1a). Using modified biochemical protocols first established by Ichihashi *et al*. (17), purified intact MVs were stripped of their membranes using detergent and reducing agents. Isolated core-LB fractions were then subjected to controlled proteolysis, using varying concentrations of trypsin, to remove LB components while minimizing core degradation. When samples were subjected to immunoblot analysis for membrane (A17L), LB (F17R) and inner core (A10L) proteins, 125ng/ml trypsin appeared to provide optimal LB removal while maintaining core integrity (Fig S1a). When carried over to large scale preparations for comparative QMS this fractionation protocol resulted in 97% membrane removal, and 98% LB removal with only a 3% loss in core integrity (Fig S1b). Electron microscopy (EM) confirmed that this protocol provides two enriched sub-virion fractions: membrane-free cores with associated LBs (LB-Core), and intact cores in which the LBs have been effectively removed (Core) (Fig S1c)

**Fig 1.**
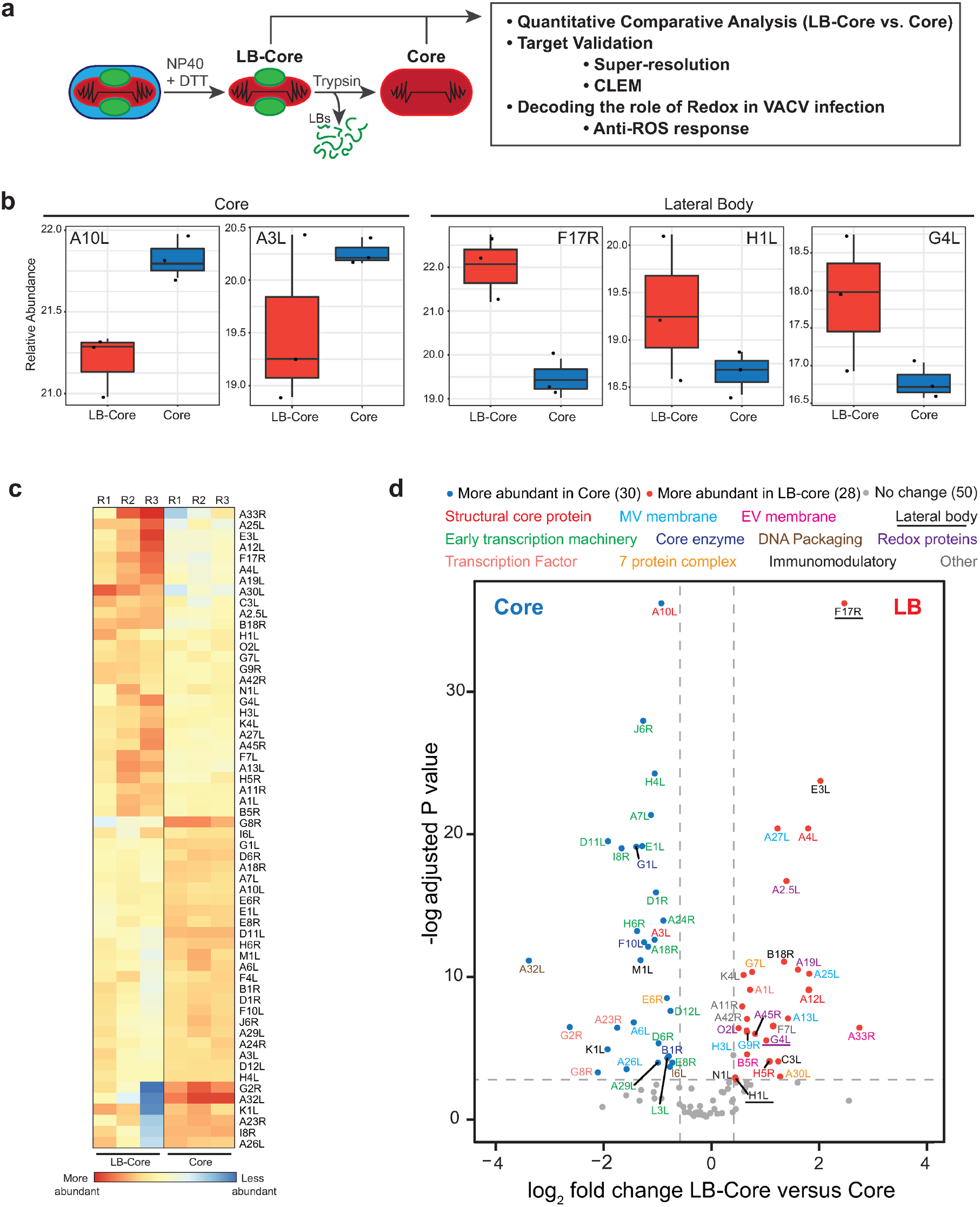
Defining the VACV LB proteome. **(a)** Schematic of controlled degradation of VACV MVs used in this study (details in Materials and Methods). **(b)** Relative abundance plots of known core (A10L, A3L) and LB (F17R, H1L, G4L) proteins across LB-Core and Core fractions. **(c)** Heat map showing the relative abundance of VACV proteins between LB-Core and Core fractions. **(d)** Relative abundance of viral proteins in LB-Core versus Core fractions. Proteins are colour coded with regard to LB-Core vs. Core abundance: more (red), less (blue) or no difference (grey). Experiments were performed in triplicate and significance scored as a minimum of two-fold greater in LB-Core vs. Core samples. An adjusted p-value ≤ 0.08 cut off was set using the adjusted p-value of the *bone fide* VACV LB protein H1L (c-d).

For spatial proteotype analysis, enriched LB-Core and Core samples were subjected to lysC-assisted tryptic digestion under pressure-cycling conditions (56, 57). Samples were analysed by data-dependent acquisition (DDA) liquid chromatography mass spectrometry (LC-MS/MS) and quantified across the two conditions. Consistent with previous reports, a total of 108 viral proteins were identified (Table S1; Tab 1). For quality control, the abundance of *bona fide* core (A10L, A3L) and LB (F17R, H1L and G4L) components were compared across samples (Fig 1b). As anticipated, the relative abundance of A10L and A3L was enriched in core samples while F17R, H1L and G4L were enriched in LB-core samples. Using the low abundance LB protein H1L as a cut off (8, 58), 58 proteins were significantly enriched in either LB-Core or Core samples (Table S1; Tab 2). Hierarchical clustering confirmed that these 58 were consistently enriched in LB-Core or Core samples across individual replicates (Fig 1c). Amongst these, 30 proteins were associated with VACV cores and 28 potential LB candidates emerged (Fig 1c, d and Table S1; Tabs 3 and 4).

Five subsets of expected core enriched proteins were identified: 2 structural core proteins (red), 15 viral early transcription machinery proteins (green), 2 DNA packaging proteins (brown), 3 core enzymes (navy), and 3 viral intermediate and late transcription factors (coral) (Fig 1c and Table S1; Tab 4). Five of the identified proteins have not been previously associated with VACV cores: VACV seven protein complex protein E6R (orange), VACV membrane proteins A26L and A6L (cyan), and the immunomodulatory proteins M1L and K1L (black) (Fig 1d).

The proteins enriched in LBs included the previously described LB proteins F17R, H1L and G4L (underlined), 6 immunomodulatory proteins (black), 5 putative redox modulating proteins (purple) and 2 seven protein complex proteins (orange) (Fig 1d, Table S1; Tab3, Table S3). We also detected 3 core proteins (red), 5 MV membrane proteins (cyan), 2 EV membrane proteins (pink), 1 late transcription factor A1L (coral), and finally 4 proteins that did not fall into a definable category (grey): 1 protein required for crescent formation (A11R), the VACV nicking/joining enzyme K4L, VACV profilin A42R and 1 protein with no defined function F7L (Table S3).

Visualizing our findings in light of the VACV MV protein-protein interaction network uncovered by Mirzakhanyan and Gershon (59), we built a schematic of our candidate LB proteins and their proposed interactions (Fig S2). The extensive connections between the identified proteins suggests that they are in close proximity within virions and explains the presence of ‘contaminating’ VACV membrane and core components within the LB proteome. Taken together, as the sub-viral localization of 13 of these proteins has been defined as MV/EV membrane, core and LB (8, 10), we were left with 15 novel LB candidate proteins (A1L, A2.5L, A11R, A19L, A30L, A42R, A45R, B18R, C3L, E3L, F7L, G7L, K4L, N1L, O2L).

### Host proteins reside in VACV LBs

As the VACV MVs used for these studies were purified from human cells, we had the opportunity to extend our analysis of the VACV LB proteome to include human proteins. Doing so, we identified a total of 586 human proteins associated with VACV virions (Table S2; Tab 1). While we confirmed 78% (60), 67% (61) and 80% (62) of the VACV-associated cellular proteins found in previous MS studies, improved MS methodology and detection allowed us to identify many more. Of the 586 cellular proteins detected, 210 were significantly enriched in LB-Core or Core samples (Table S2; Tab2), while the remaining 376 proteins were not enriched in either. Hierarchical clustering across individual replicates showed that amongst the 210 enriched proteins, 93 were consistently found in LB-Core samples and 117 in Core samples (Fig 2a & b and Table S2; Tabs 3 and 4). In the comparative quantitative analysis, 6 human proteins (mitochondrial transcription factor A, vimentin, Ankyrin repeat domain-containing protein 17, NADH dehydrogenase 1 alpha subcomplex subunit 7, X chromosome RNA-binding motif protein, mitochondrial 39S ribosomal protein L14) were more significantly enriched at a greater abundance than the major LB viral protein F17R, suggesting that these human proteins are major LB constituents.

**Fig 2.**
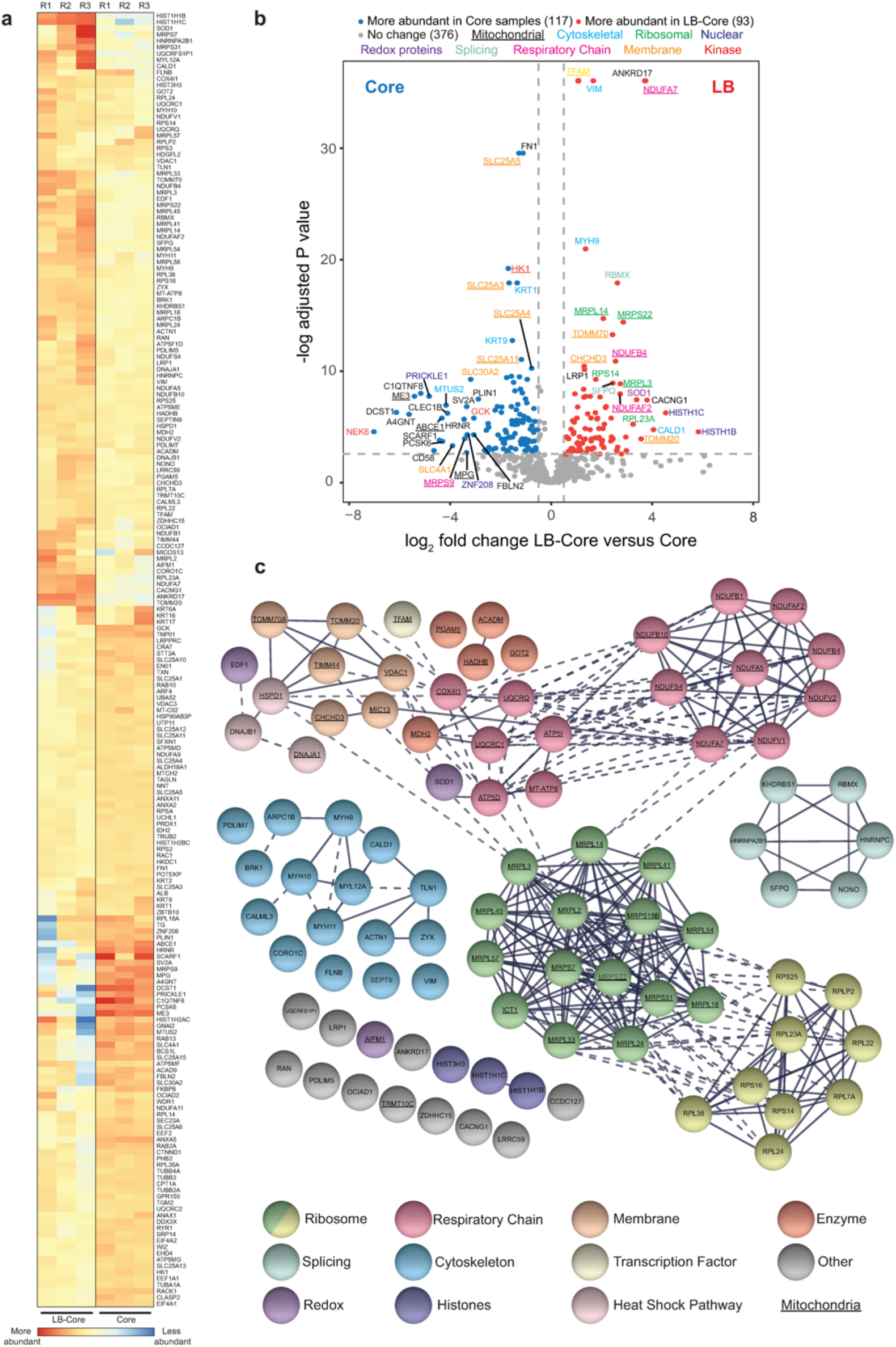
Cellular protein constituents of VACV lateral bodies. **(a)** Heat map showing the relative abundance of human proteins in LB-Core vs. Core fractions. **(b)** Relative abundance of human proteins in LB-Core vs. Core fractions. Proteins are colour coded with regard to LB-Core vs. Core abundance: more (red), less (blue) or no difference (grey). Experiments were performed in triplicate and significance scored as a minimum of two-fold for LB-Core relative to Core with an adjusted p value ≤ 0.08 (a & b). **(c)** STRING protein-protein interaction network of human proteins enriched in LB-Core vs. Core fractions. Proteins were clustered and coloured according to function as indicated. Proteins involved in mitochondrial function are underlined. Solid lines depict confirmed physical interaction (confidence >0.9), dashed line indicates suggested co-location in screening studies.

String visualization of the human proteins within LBs revealed 7 major protein interaction clusters (Fig 2c and Table S2; Tab 3) (63). These included 9 ribosome subunits, 6 proteins involved in splicing and 16 cytoskeletal-associated proteins including regulators of actin dynamics, myosins and intermediate filaments (Fig 2 and Table S2; Tab 3). Strikingly, over half (45) of the candidate human LB proteins were of mitochondrial origin including 15 proteins of the respiratory chain, 15 mitochondrial ribosomal subunits, 5 mitochondrial enzymes and 6 proteins involved in mitochondrial membrane stability and import.

### VACV LBs harbour a set of viral proteins with redox modulating potential

The presence of so many mitochondrial proteins in LBs along with the co-identification of five putative viral redox proteins (G4L, A2.5L, A19L, A45R and O2L) prompted us to focus on these LB candidates. Briefly, with the exception of A45L, which is 39% identical to human Cu-Zn SOD (51), each of these proteins contain Cxx(x)C motifs characteristic of redox modulating enzymes (64). Amongst these, the glutaredoxin G4L, a homologue of human Glutaredoxin-2, is the only previously identified LB protein (8). Together with A2.5L, it has been reported to be part of a VACV encoded cytoplasmic disulphide formation pathway that is essential for virus assembly (30–32, 52). The VACV protein A19L carries two CxxC motifs that are important for interaction with VACV core transcription machinery and are essential for VACV infection (53). A45R is a non-functional SOD-like dismutase, whose *Leporipoxvirus* homologue has been shown to disrupt host SOD1 activity (51, 65, 66), and finally, the O2L protein is a homologue of human Glutaredoxin-1 that contains a single CxxC motif and displays *in vitro* thioltransferase activity (50, 55, 67).

Relative abundance plots of the MS data indicated that G4L, A2.5L, A19L, A45R and O2L were all significantly enriched in LB-core over core samples (Fig 3a). While indicative of LB residence, to validate the MS results and confirm the localization of these proteins to LBs we turned to super-resolution microscopy. We have previously used structured illumination microscopy (SIM), stimulated emission depletion (STED), and stochastic optical reconstruction microscopy (STORM), in combination with single particle averaging to map viral proteins to distinct VACV substructures including LBs, cores and the viral membrane (68–72).

**Fig 3.**
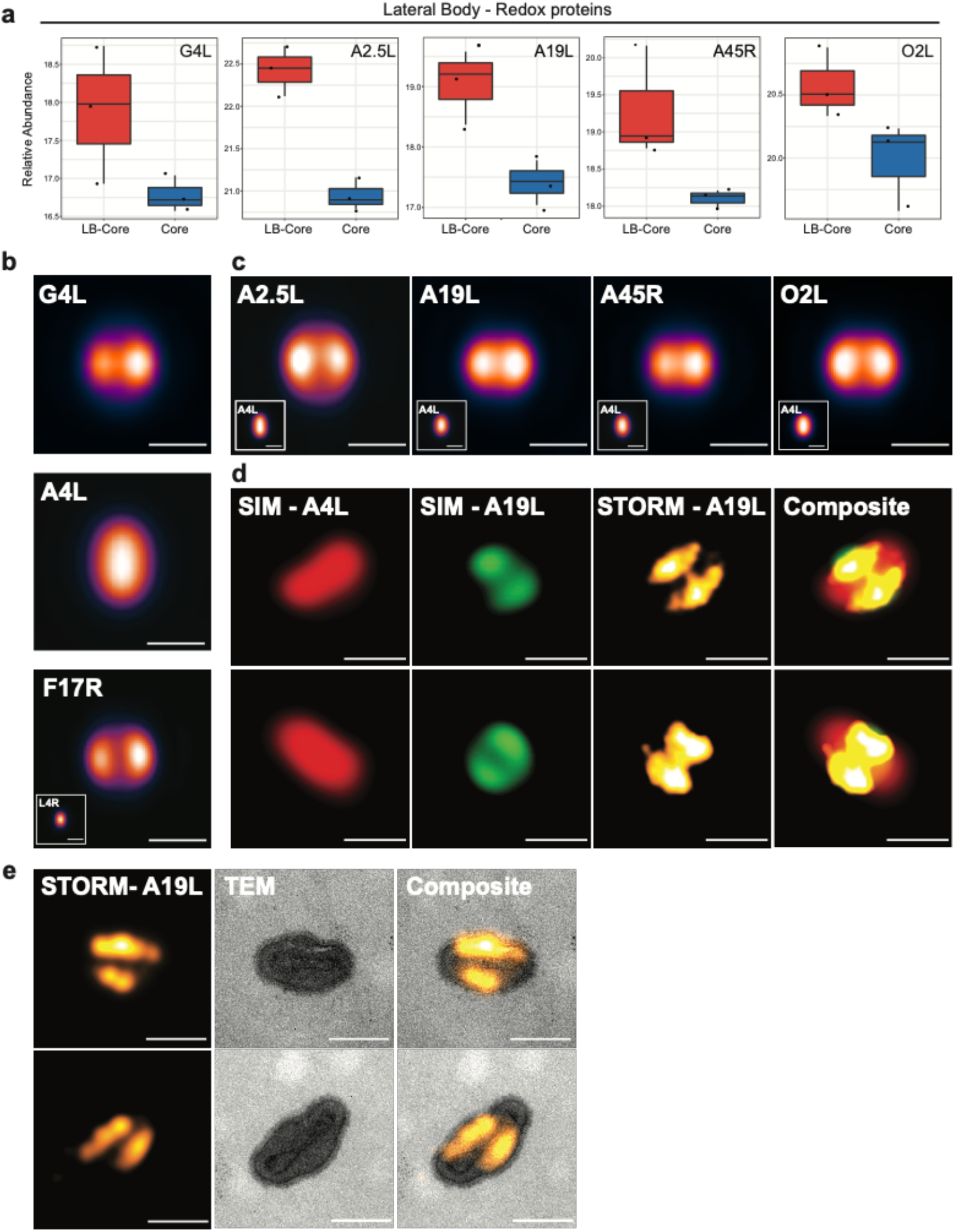
Viral redox proteins reside in LBs. **(a)** Relative abundance plots of the five putative redox modulating viral enzymes (G4L, A2.5L, A19L, A45R, O2L) across LB-Core and Core fractions **(b)** VirusMapper models (sagittal orientation) of G4L (LB) and A4L (core) protein localization using WR mCherry-A4L G4L-EGFP virions (n= 949). A VirusMapper model of WR L4R-mCherry EGFP-F17R is displayed for comparison (n= 231) **(c)** VirusMapper models (sagittal orientation) of EGFP-tagged redox proteins: A2.5L, A19L, A45R, O2L (n > 100). **(d)** Correlative SIM-STORM imaging of WR mCherry-A4L A19L-EGFP virions. Two representative virions are displayed (Overview in Fig S3a) **(e)** Correlative STORM-TEM of WR mCherry-A4L A19L-EGFP virions immunolabelled with anti-GFP nanobody directly conjugated to AlexaFluor647-NHS. STORM images of A19L were registered with EM micrographs. Two representative virions are displayed (Overview Fig S3b). **(b-e)** Scale bars = 200 nm.

To this end, we built a library of recombinant VACVs using the strategy employed for the successful identification of the major LB protein F17R (8). A parental virus harbouring a mCherry-tagged version of the core protein A4L was used to generate a set of recombinant viruses that carry an additional EGFP-tagged version of G4L, A2.5L, A19L, A45R or O2L Having shown that G4L is a LB protein using immuno-EM (8), we used it as a test case. Virions were imaged by dual-colour SIM and localization models of G4L and A4L were generated as previously described (68, 71). Using A4L to identify virion position and orientation (Fig 3b; middle panel), the localization models showed that G4 resides in two elliptical structures at the sides of cores corresponding to VACV LBs (Fig 3b; top panel). A model of virions containing EGFP-tagged F17R, the major *bone fide* LB protein, is displayed for comparison (Fig 3b; bottom panel).

Applying this imaging and analysis pipeline to the remaining VACV redox proteins we confirmed, consistent with the MS results, that A2.5L, A19L, A45R and O2L are each novel VACV LB proteins (Fig 3c).

To further exemplify the utility of this super-resolution virion protein mapping approach, we performed correlative SIM-STORM and correlative STORM-TEM (transmission electron microscopy) on mCherry-A4 A19L-EGFP recombinant VACV. For both, virions were permeabilised and immuno-labelled with AlexaFluor647-NHS conjugated anti-GFP nanobodies to enable STORM imaging of the A19L-EGFP. For SIM-STORM, virions were imaged using dual-colour SIM for core (A4L) and LB (A19L), followed by far-red STORM reporting on the fluorescently-labeled nanobodies. Representative virions displayed in Figure 3d (overview Fig S3a), show VACV cores (SIM-A4L) flanked by two A19L positive LBs (SIM-A19L). As anticipated, the A19L STORM reconstructions largely overlap with the SIM A19L images, albeit with better resolution (Fig 3d; STORM-A19L). For STORM-TEM, virions were first imaged by STORM for A19L-EGFP, followed by EM sample preparation, sectioning and imaging of the same virions by TEM. Correlation of these images supports the SIM and STORM data, confirming that A19L resides in VACV LBs (Fig 3e; overview Fig S3b).

Together these results serve to confirm our mass spectrometry data, which identifies viral redox proteins as a new class of VACV LB constituents and demonstrate the utility of correlative super-resolution imaging to define the sub-viral location of proteins in poxviruses.

### VACV infection exerts an anti-oxidative effect on host cells

All virus infections exert a redox imbalance in the host cell and intracellular RONS production. This in turn activates anti-viral immune responses such as the upregulation of autophagy, promotion of cell death, inhibition of mTOR and Pattern Recognition Receptor downstream signalling events (36). Given the presence of five putative redox modulating enzymes in LBs we considered the possibility that VACV modulates cellular RONS production for successful infection. To assess this, we first looked at the impact of VACV infection on ROS levels generated during normal cellular respiration. Cells were infected with WT VACV and the percentage of ROS positive cells quantified at 1 h and 8 h post infection (pi) using an oxidant-sensitive probe in combination with flow cytometry. These time points correspond to early infection, when LB proteins are delivered into the cytoplasm by incoming virions, and late infection when greater amounts of these LB proteins are newly synthesized. Untreated, uninfected cells served as the negative control and uninfected cells treated with tert-butyl hydroperoxide (TBHP), a ROS inducer, were included as a positive control for ROS activation. At both time points ~40% of untreated, uninfected control cells were ROS positive (Fig 4a). Infection with VACV resulted in a >3-fold reduction in ROS positive cells at h pi (12%), and a >23-fold reduction by 8 h pi (1.7%) (Fig 4a).

**Fig 4.**
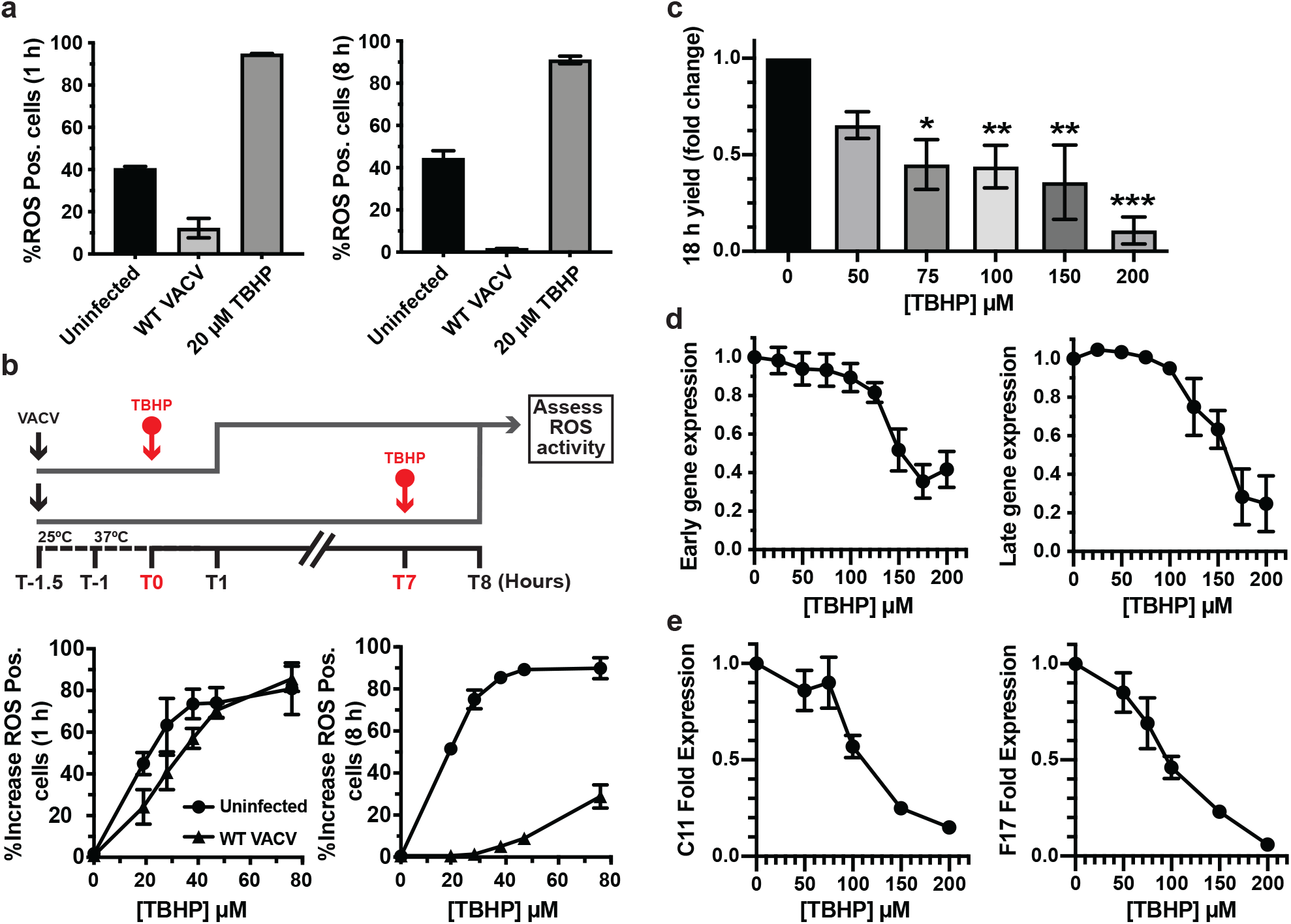
VACV supresses host cell oxidative responses to promote infection. **(a)** The percentage of ROS activated A549 cells measured at 1 and 8 h pi in cells left uninfected (black bar) or infected with WT VACV (Light grey bar). As a positive control, uninfected cells were treated with 20 μM TBHP for the final hour of the experiment (dark grey bars). **(b)** Top: schematic of experiment to assess impact of VACV infection on ROS activation. VACV was bound to A549 cells for 30 min at RT prior to being shifted to 37 °C for 1 h. Cells were challenged with the indicated concentrations of ROS inducer TBHP for 1 h prior to quantification of ROS-positive cells at 1 h pi and 8 h pi. Bottom: The percentage of uninfected (circles) and infected (triangles) ROS positive cells at 1 h pi and 8 h pi challenged for the final hour with the indicated concentrations of TBHP (n = 3; ± SEM). **(c)** A549 cells were infected with WT WR VACV (MOI=1) in a range of TBHP concentrations. At 18 h pi, the infectious virus yield was assessed by plaque assay (n=3; mean ± SEM). **(d)** A549 cells infected with WR E-EGFP or L-EGFP were challenged with the indicated concentrations of TBHP for 8 h. Early and late viral gene expression was assayed by flow cytometry and the data plotted as the fold change from infected untreated cells (n=3; mean ± SEM). **(e)** A549 cells infected with WT VACV were challenged with the indicated concentrations of TBHP for 8 h. Total RNA was extracted and early (C11R) and late (F17R) gene expression assessed by RT-qPCR. Data is displayed as fold expression normalised to GAPDH. (n = 3 ± SEM, *P<0.05, **P<0.005, ***P<0.0005)

To test how effective VACV is at shunting ROS activation, cells infected with WT VACV were challenged with increasing concentrations of TBHP at 0 h or 7 h pi and the increase in ROS positive cells determined at 1h and 8 h pi (Illustrated in Fig 4b; top). Compared to uninfected cells, at 1 h pi VACV infection reduced the number of ROS positive cells by 1.84-fold at low TBHP concentrations prior to being overcome at concentrations ≥ 50 μM (Fig 4b; lower left). At 8 h pi post infection, VACV infection potently attenuated ROS activation by TBHP (10-fold at 50 μM relative to uninfected controls, with a 28.8% break through at 80 μM) (Fig 4b, lower right). These results indicated that VACV exerts an anti-oxidant effect on host cells both early and late during infection, a finding consistent with LB-mediated delivery of redox proteins and their subsequent post-replicative expression.

### High levels of ROS are detrimental to VACV infection

Despite the fact that VACV encodes and packages 5 proteins with redox modulating potential, we found that the artificial induction of high ROS levels could not be fully supressed during infection. This prompted us to ask if elevated cellular ROS levels would be deleterious to VACV replication. Cells were infected with WT VACV in the presence of various concentrations of TBHP (50 μM-200 μM) and the infectious yield determined at 18 h pi. We observed a dose-dependent effect with a 1.5-fold reduction in virus yield at 50 μM TBHP, and a 3.7-fold reduction at 200 μM TBHP (Fig 4c). LDH cytotoxicity assays of cells treated with the same concentrations of TBHP indicated that this reduction was not due to cell death (Fig S4a). To determine at what stage of VACV infection ROS-mediated inhibition occurred we measured VACV early and late gene expression using recombinant viruses that express EGFP under the control of endogenous early (WR E-EGFP) or late (WR L-EGFP) viral promoters. Increasing concentrations of TBHP resulted in decreasing numbers of cells expressing early and late genes as determined by flow cytometry at 8 h pi (Fig 4d). LDH assays confirmed that this decrease was not due to cell death (Fig S4b). To further define whether this block in viral gene expression was at the level of transcription or translation, cells were infected with WT VACV in the presence of increasing concentrations of TBHP. At 8 h pi RT-qPCR was used to determine the vRNA levels of the VACV early gene (C11L) and a VACV late gene (F17R) (73). A dose-dependent inhibition of both early and late viral gene transcription was seen with increasing TBHP concentrations (Fig 4e). This indicated that the block in VACV gene expression seen upon increasing oxidative stress occurs at the level of transcription and serves to effectively shunt the expression of early viral genes.

Collectively these results demonstrate that VACV infection actively and effectively down regulates the cellular ROS production, and that conversely high levels of ROS have a deleterious effect on VACV transcription, protein expression and subsequent virion production.

## Discussion

All members of the *Poxviridae* carry LBs between their membrane and core (19). Using EM, Dales first observed and described LBs as ‘lateral dense elements’ which disassociate from the virus core during entry (74). *In vitro* experiments on purified VACV virions soon followed, showing that LBs remain associated with viral cores upon biochemical removal of the virion membrane (18). Together these findings suggested the poxviruses were composed of multiple structural elements and hinted that LBs were independent entities. *In vitro* studies by Ichihashi and Oie confirmed that LBs were distinct from virus cores and composed of proteins (17).

Using similar techniques, we identified the first three *bona fide* viral LB proteins (8): F17, a small basic VACV phosphoprotein, the VACV phosphatase H1, and G4, a viral glutaredoxin required for cytoplasmic disulphide formation during virion assembly (30). We showed that F17 was the major LB protein (69% of LB mass) and that it is degraded in a proteosome-dependent manner to release H1 (1% of LB mass) during deposition of virus cores into the cytoplasm. Upon release H1 fulfils its effector function - blocking STAT1 nuclear translocation and preventing STAT1-mediated anti-viral responses - prior to early gene expression. Based on these findings we speculated that LBs were akin to Herpesvirus tegument, serving to deliver immune- or cell-modulating proteins into host cells, allowing the virus to establish its replicative niche (8, 75).

While LBs appear to play an essential role in the poxvirus lifecycle, the composition of LBs and the function of their constituents has never been fully defined. This is partially due to difficulties associated with isolation or purification of LBs from cells or virions, and that the vast majority of studies are from the pre-omics era. Taking advantage of spatial proteotyping, we could assess protein enrichment across biochemically fractionated virion samples, thereby circumventing the need for LB isolation or purification (17, 18).

This approach also provided us with the opportunity to ask, for the first time, if host proteins are packaged into VACV LBs. To this end, we identified 93 human proteins enriched in LBs. The majority of proteins identified were cytoskeletal, ribosomal or mitochondrial in origin. While their presence could be due to erroneous packaging during cytoplasmic assembly, we speculate that VACV packages and delivers these factors to facilitate establishment of its replicative niche. Strikingly, 6 of the host proteins were found to be more abundant in LBs than F17 raising the possibility of active LB incorporation. One of these host proteins, vimentin, is known to localize to viral replication sites (76) and is packaged into virions (61, 77) where it, like F17, biochemically fractionates with Core-LBs and is trypsin sensitive (78). Further confirmation and research into these host factors would be highly valuable.

While this is the first study to look at host proteins in VACV LBs, there are two previous reports using similar fractionation/degradation protocols to identify VACV structural components (17, 79). Though well performed, due to technical limitations at the time, one study lacks genomic information and the other is limited to N-terminal sequencing. This makes it difficult to compare the findings between one another or with our own. Nonetheless of the 6 viral proteins identified in the latter study; 3 described as core / core-LB are *bona fide* core proteins (A3, A10, A12), 2 described as membrane-LB are *bona fide* membrane proteins (A17, D8) and the remaining protein described as core-LB (G7) we identified in this study as a LB component. As a member of the VACV 7-protein complex, G7 is a required for the association of viral membrane with the to-be intra-virion components during VACV assembly (80). As postulated by Condit and Moussatche (81, 82), consistent with the presence of F17 in LBs we also identified A30, a 7-protein complex member that interacts with both F17 and G7 (83). Our finding that the other 7-protein complex members including, the F10 kinase, were not identified as LB components is consistent with the previously described sub-viral compartmentalization of viral phosho-enzymes during virion maturation (84). It will be of future interest to determine if, in addition to their role in virion assembly, F17, G7 and A30 act as LB-delivered effectors in any capacity.

In this report we focused on confirming the LB localisation of five viral proteins with redox modulating capabilities; A2.5L, A19L, A45R, O2L and the previously identified G4 (8). Super-resolution imaging was used to unequivocally show that these proteins are LB components. Together with the fact that VACV replicates exclusively in the cytoplasm, this strongly suggested that controlling the production of intracellular ROS is important for VACV replication. Consistent with LB delivery of redox modulators, VACV infection diminished basal cellular ROS levels as early as 1 h pi. We have previously shown that the reducing environment of the cytoplasm is essential for the dismantling of incoming disulphide-linked core proteins to allow for core expansion and subsequent early gene transcription (8). Consistent with this, we show here that exogenous activation of ROS is detrimental to VACV infection at the level of early gene transcription. Collectively this data suggests that, like cytomegalovirus, VACV has evolved to overcome the anti-viral oxidative challenge presented by cells during infection (46, 47). In support of this, IFNγ-driven NO production inhibits VACV and ectromelia virus replication in mouse macrophages (48), and NOS2-deficient mice are highly susceptible to ectromelia despite no known loss of ectromelia-targeted immune responses (49). Taken together with our findings, these results suggest that poxviruses must overcome RONS for successful replication.

Previous work indicates that A45 and O2 are not required for VACV replication in tissue culture, that A45 is dispensable *in vivo* and that A2.5, G4 and A19 are each required for virus assembly (30–32, 50–55). While this built-in functional redundancy and dual functionality makes investigating the impact of the individual factors challenging, future experimentation will be aimed at confirming the role of the individual VACV redox proteins in overcoming cellular RONS and extending this to mechanistic understanding. Akin to the VACV strategy of dedicating 9 different yet redundant viral factors to inhibit the NF-kB pathway (85, 86), we postulate that the various redox proteins target RONS signalling through divergent immunomodulatory targets.

In summary, by combining comparative quantitative mass spectrometry, super-resolution microscopy and molecular biology we defined the poxvirus LB proteome. Furthermore, we uncovered a correlation between the packaging of viral redox proteins in LBs, the ability of VACV to disrupt the intracellular oxidative environment and the inhibition of VACV infection by ROS. The presence of all VACV-encoded redox proteins in LBs underscores the importance of blocking RONS for viral replication. In addition to defining the LB proteome as a valuable resource to the fields of virology and immunology, this study provides a solid foundation for mechanistic studies aimed at understanding the role of RONS in the cellular control of poxvirus infection and consequently how viruses have evolved to evade this.

## Materials and methods

### Cell lines and virus propagation

BSC40 (kind gift of P. Traktman, Medical University of South Carolina, Charleston, SC, USA), hTert RPE-1 (Clonetech laboratories), A549 (ATCC CCL-185) and HeLa cells (ATCC CCL-2) were grown in Dulbecco’s modified Eagle’s medium (DMEM) supplemented with 100 units/ml penicillin, 100 μg/ml streptomycin, 1 mM sodium pyruvate, 100 μM non-essential amino acids and 10 % heat inactivated FCS. A549 cells were seeded onto fibronectin coated plates for all assays. VACV MVs were propagated in BSC40 or RPE cells and purified from cell lysates through a sucrose cushion prior to band purification on a sucrose gradient, as described previously (20). All viruses used in this study were based on VACV strain Western Reserve. WR E-EGFP, WR L-EGFP, WR mCherry-A4L and WR L4R-mCherry EGFP-F17R [WR VP8-mCherry EGFP-F17] have been previously described (8, 87, 88).

### Viral Fractionation

MVs were fractionated into sub-viral constituents by incubation with 1% NP-40 and 50 mM DTT in 10 mM Tris, pH 9.0, for 30 min, 37 °C with gentle perturbation. The insoluble fraction was sedimented for 1 hr at 21,130 x g, 4 °C and the supernatant (Sup 1; Membrane fraction) was retained. The insoluble fraction was washed (Wash 1) by resuspension in 10 mM Tris pH 9.0 and sedimented for 1 hr at 21,130 x g at 4 °C (Pellet 1; LB-Core). Both fractions were retained for analysis. For LB removal, LB-Core fractions were treated with trypsin (31.25, 62.5, 125, 250, 500, 1000 ng/ml) as stated, in 10 mM Tris pH 9.0 for 15 min, 37 °C prior to addition of 400 μg/ml Soybean Trypsin Inhibitor. The insoluble fraction was sedimented 1 hr at 21,130 x g at 4 °C (Pellet 2; Core) and, along with the soluble supernatant fraction (Sup 2), retained for analysis. For subsequent fractionations prior to MS or EM analysis, 125 ng/ml Trypsin was applied, and an additional wash performed.

### Electron Microscopy (EM)

LB-Core and Core samples were prepared as described (see <u>Viral Fractionation)</u> and prepared for EM by fixation in 1.5% glutaraldehyde / 2% EM-grade paraformaldehyde (TAAB) in 0.1M sodium cacodylate for 45 min at RT. Briefly, samples were treated with reduced osmium, tannic acid, dehydrated through an ethanol series and embedded in epon resin (89). Sections were collected on formvar coated slot grids, stained with lead citrate and imaged using a transmission electron microscope (Tecnai T12, Thermo Fisher Scientific) equipped with a charge-coupled device camera (SIS Morada; Olympus).

### Immunoblotting

Immunoblot samples were boiled in LDS DTT sample buffer (Novex, Life Technologies) for 5 min at 98 °C. Proteins were separated on 4-12% or 12% Bis-Tris gels (Invitrogen), transferred to nitrocellulose membranes, and analysed using rabbit anti-A10L (1:1,000), anti-F17R (1:1,000), anti-A17L (1:500) in combination with horseradish peroxidase (HRP)-coupled secondary antibodies (1:2,000) and Immoblion Forte Western HRP substrate (Merck) or with IR-Dye secondary antibodies (Licor) (1:10,000). Blots were visualised using photographic film and X-ray developer or LiCor Odyssey. Relative quantification of immunoblots was performed using Fiji.

### Mass Spectrometry (MS)

Lysis and tryptic digestions were performed under pressure-cycling conditions using Barocycler NEP2320-45k (PressureBioSciences, South Easton, MA) as described previously (56). Briefly, LB-Core and Core fractions were resuspended in 8 M urea containing 100 mM ammonium bicarbonate pH 8.2, 10% 2,2,2 trifluoroethanol, one tablet of phosphatase inhibitors cocktail (PhosStop, Roche) per 10 ml buffer. Lysis was performed under pressure cycling conditions (10s at 45 kpsi, 10s at 0 Kpsi, 297 cycles). Samples were sonicated three times for 30s and then rested for 1 min on ice and spun at full-speed in a bench top centrifuge for 5 min. Supernatants were treated with 10 mM TCEP for 20 min at 35 °C and then 40 mM iodoacetamide in the dark at RT for 30 min. Samples were diluted to contain 6 M urea and digested with LysC 1/50 w/w under pressure cycling conditions (25s at 25 Kpsi, 10s at 0 Kpsi, 75 cycles). Samples were further diluted to 1.5 M urea and digested with 1/30 w/w trypsin was performed pressure cycling conditions (25s at 20 Kpsi, 10s at 0 Kpsi, 198 cycles). Samples were mixed by inversion overnight at 37 °C and digestion stopped by the addition of trifluoroacetic acid to pH 2.0. Resultant peptides were desalted on a reverse phase C18 column (Waters corp., Milford, MA) and eluted with 40% acetonitrile (ACN), 0.1% TFA. The solvents were evaporated using a centrifuge evaporator device. Peptides from LB-Core and Core samples were resuspended in 2% ACN, 0.1 % formic acid subjected to direct LC-MS/MS analysis.

Peptide samples from LB-Core and Core samples were separated by reversed-phase chromatography on an ultra-high-pressure liquid chromatography (uHPLC) column (75 μm inner diameter, 15 cm, C18, 100 Å, 1.9 μm, Dr Maisch, packed in-house) and connected to a nano-flow uHPLC combined with an autosampler (EASY-nLC 1000, Thermo Scientific). The uHPLC was coupled to a Q-Exactive Plus mass spectrometer (Thermo Scientific) equipped with a nanoelectrospray ion source (NanoFlex, Thermo Scientific). Peptides were loaded onto the column with buffer A (99.9% H_2_O, 0.1% FA) and eluted at a constant flow rate of 300 nl per min over a 90 min linear gradient from 7% to 35% buffer B (99.9% ACN, 0.1% FA). After the gradient, the column was washed with 80% buffer B and re-equilibrated with buffer A. Mass spectra were acquired in a data-dependent manner, with an automatic switch between survey MS scan and MS/MS scans. Survey scans were acquired (70,000 resolution at 200 m/z, AGC target value 10^6^) to monitor peptide ions in the mass range of 350–1,500 m/z, followed by higher energy collisional dissociation MS/MS scans (17,500 resolution at 200 m/z, minimum signal threshold 420, AGC target value 5 × 10^4^, isolation width 1.4 m/z) of the ten most intense precursor ions. To avoid multiple scans of dominant ions, the precursor ion masses of scanned ions were dynamically excluded from MS/MS analysis for 10 s. Singly charged ions and ions with unassigned charge states were excluded from MS/MS fragmentation.

COMET (release 2015.01 rev. 2) (90) was used to search fragment ion spectra for a match to tryptic peptides allowing up-to two missed cleavage sites from a protein database, which was composed of human proteins (SwissProt, v57.15), Vaccinia virus proteins (UniProt, strain Western Reserve, v57.15), various common contaminants, as well as sequence-reversed decoy proteins. The precursor ion mass tolerance was set to 20 ppm. Carbamidomethylation was set as a fixed modification on all cysteines. The PeptideProphet and the ProteinProphet tools of the Trans-Proteomic Pipeline (TPP v4.6.2) (91) were used for probabilistic scoring of peptide-spectrum matches and protein inference. Protein identifications were filtered to reach an estimated false-discovery rate of ≤1%. Peptide feature intensities were extracted using the Progenesis (v2.0) LC-MS software (Nonlinear Dynamics). Protein fold changes and their statistical significance between paired conditions were tested using at least two fully tryptic peptides per protein with the MSstats library (v1.0) (92). Resulting p-values were corrected for multiple testing with Benjamini-Hochberg method (93).

### STRING Analysis

The LB human protein-protein interaction network analysis was performed using the STRING database at high stringency (0.700), clusters were generated using kmeans clustering (8) (https://string-db.org).

### Generation of recombinant vaccinia viruses

Novel recombinant viruses were generated by homologous recombination as previously described (20). Recombinant dual fluorescent viruses were generated using plasmids based on pJS4 for insertion of a second copy of the LB candidate protein with a C-terminal EGFP fusion protein into the Tk locus of WR mCherry-A4 (94). For the G4L dual fluorescent virus, the EGFP fusion was N-terminal.

### Flow cytometry

Cells were detached with 0.05 % trypsin-EDTA, trypsin inactivated by addition of 5% FBS and fixed with 1% Formaldehyde in PBS. Samples were analysed for EGFP using a Guava easyCyte HT flow cytometer and InCyte software (Millipore).

### Super-Resolution Imaging and Correlative light and electron microscopy

Band purified MVs diluted in 20 μl 1 mM Tris pH 9.0 were sonicated and vortexed prior to being placed on coverslips for 30 min. Bound virus was fixed with 4% formaldehyde. When required, samples were permeabilized for 30 min with 1% TritonX-100 in PBS and blocked with 5% BSA (Sigma), 1% FCS for 30 min and immune-stained with anti-GFP nanobody (Chromotek) conjugated in-house to AlexaFluor647-NHS (Invitrogen) in 5% BSA, PBS at 4 °C overnight. Coverslips were washed three times with PBS prior to mounting. Both mounting and imaging by STORM and SIM and Virusmapper modelling have been described previously (68). Correlative super-resolution light and electron microscopy imaging of MVs was performed as previously described (95).

### Viral Yield Assay

A549 cells were infected with WT VACV (MOI 1) for 1 h. Cells were treatment with 0-200 μM TBHP as indicated and incubated at 37 °C, 5 % CO_2_ for 18 h. Cells were collected and resuspended in 1 mM Tris pH 9.0. MVs were titred on BSC40s. Forty-eight h pi, cells were fixed, and viral plaques visualized using 0.1 % crystal violet/ 2 % formaldehyde. Titres were normalised to 0 μM TBHP condition.

### Cytotoxicity assay

A549 cells were treated with 0-200 μM TBHP for 8 h or 18 h. Cytotoxicity was measured using the Pierce LDH Cytotoxicity Assay Kit (ThermoScientific) according to the manufacturer’s instructions. Absorbance was measured at 490 nm and 650 nm using a VERSA max microplate reader (Molecular Devices).

### Viral gene expression assays

For flow cytometry experiments, A549 cells were infected with WR E-EGFP (MOI 1) or with WR L-EGFP (MOI 3). At 1 h pi cells were treated with 0-200 μM TBHP for 8 h followed by flow cytometry analysis.

### Reverse transcription – quantitative PCR (RT-qPCR)

For RT qPCR experiments, A549 cells were infected with WT VACV (MOI 3) for 1h and cells treated with 0, 100, 200 μM TBHP for 8h followed by total cell RNA extraction. Total RNA was extracted using the RNeasy Plus Mini kit (Qiagen) according to manufacturer’s instructions. Single stranded cDNA was reverse transcribed from 500 ng of total RNA with SuperScript II Reverse Transcriptase (Invitrogen) and oligo(dT) primers (Invitrogen). cDNA was diluted 1:5 in water and μl used as template for qPCR using Mesa blue MasterMix Plus for SYBR (Eurogentec) and a CFXConnect Real-Time System (BioRad) PCR machine. The expression levels of the viral genes C11L (early), F17R (late), and the human gene GAPDH were measured using specific primers (Integrated DNA Technologies): C11L (5’-AAACACACACTGAGAAACAGCATAAA-3’ and 5’-ACTATCGGCGAATGATCTGATTA-3’), F17R (5’-ATTCTCATTTTGCATCTGCTC-3’ and 5’-AGCTACATTATCGCGATTAGC-3’) GAPDH (5’-AAGGTCGGAGTCAACGGATTTGGT-3’ and 5’-ACAAAGTGGTCGTTGAGGGCAATG-3’). Gene expression was normalised to GAPDH and then to the untreated condition for each gene.

### Fluorogenic detection of Oxidative stress

VACV MVs (MOI 50) were bound to A549 cells for 30 min at room temperature. Cells were shifted to 37 °C for 1 h. Cells were further incubated for 1 h or 8 h. Samples were treated with 250 nM CellROX green (Thermo-Fischer Scientific) for the final 30 min prior to analysis by flow cytometry. Where stated, cells were challenged with 0-80 μM TBHP for 1 h prior to flow cytometry analysis.

## Acknowledgements

We thank V. Edwards and H. Mok for initial work on the project, V. Gould for her technical support throughout the project, P. Pereira and R. Henriques for advice on super-resolution imaging. We thank all members of the Mercer lab for helpful discussions and comments on the manuscript. This work was funded by a Sir Henry Wellcome Post-doctoral Fellowship (WT106080/Z/14/Z; to SRB), MRC Programme grant (MC_UU_00012/7; to JM), the MRC (MR/ K015826/1; to JM), the European Research Council (649101, UbiProPox; to JM), and the Swiss National Science Foundation (grant 31003A_160259; to B.W.) and a Marie Skłodowska-Curie fellowship funded by the European Union (750673; to DA). JJB is supported by core funding to the LMCB (MC_U12266B).

## Supplementary Figures

**Supplementary Fig 1.**
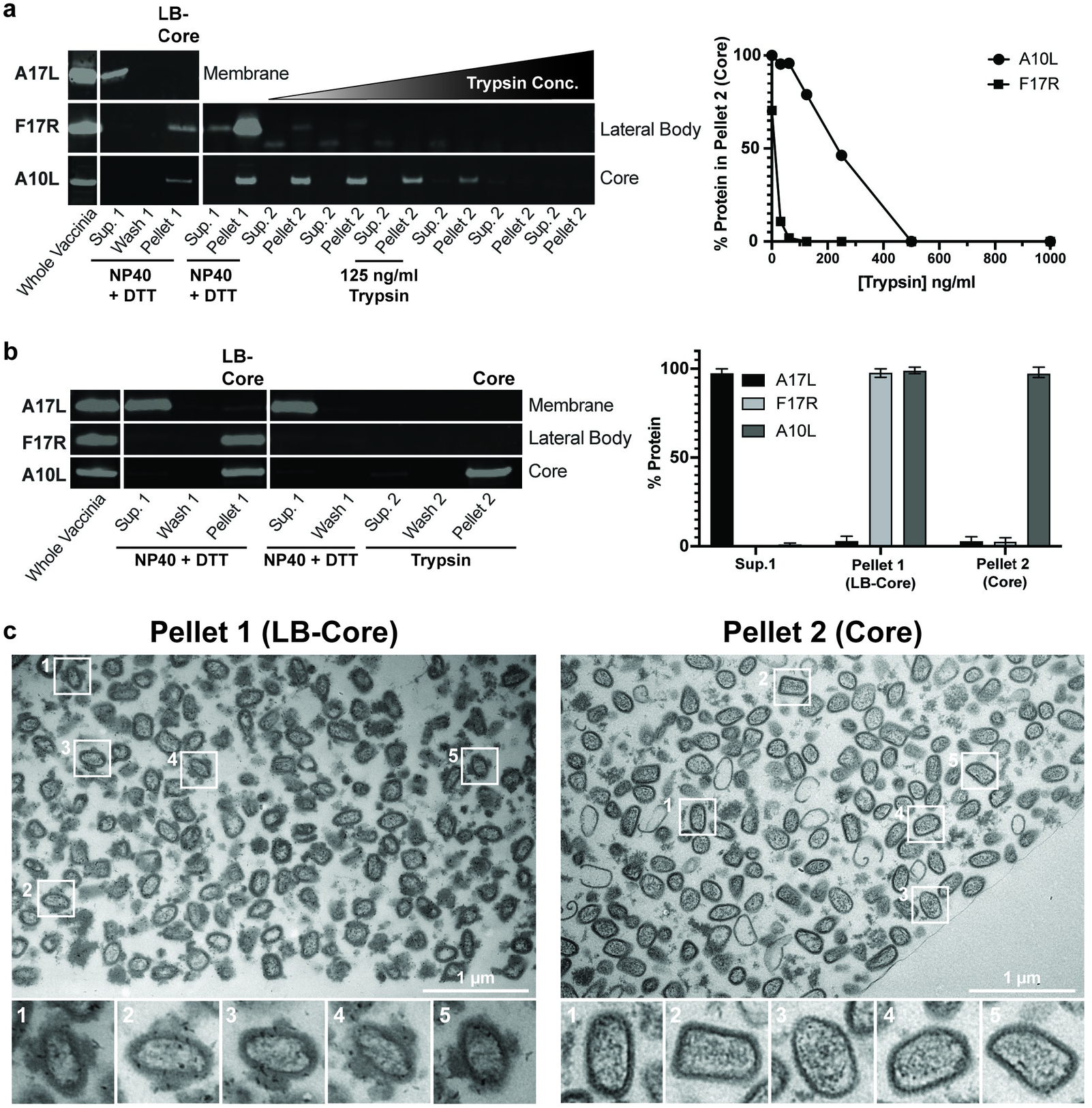
VACV Fractionation quality control. **(a)** Left: VACV MV fractionation and LB digestion conditions were optimised using WT VACV MVs. Virion membranes (Sup. 1) were removed by treatment with NP-40 + 50 mM DTT. To assess optimal conditions for LB removal, LB-Core samples (Pellet 1) were treated with various trypsin concentrations (31.25, 62.5, 125, 250, 500, 1000 ng/ml). The corresponding soluble (Sup. 2) and insoluble Core samples (Pellet 2) were retained. Viral fraction samples were analysed by immunoblotting against A17L (membrane protein), F17R (LB protein) and A10L (core protein). Right: the percentage of A10L and F17R residing in Pellet 2 (Core) upon increasing trypsin concentration was quantified. A trypsin concentration of 125 ng/ml was selected for preparing mass spectrometry samples **(b)** Left: Representative immunoblot of A17L (membrane protein), F17R (LB protein) and A10L (core) from a large-scale fractionation performed, as determined in (a), for mass spectrometry analysis. Right: Quantification of the percentage of A17L, F17R and A10L in Sup. 1 (Membrane), Pellet 1 (LB-Core) and Pellet 2 (Core) samples across triplicate large-scale fractionations prepared for mass spectrometry. **(c)** WT VACV MVs were fractionated as determined in (a). Pellet 1 (LB-Core) and Pellet 2 (Core) samples were washed after the trypsin treatment, fixed, cryosectioned and imaged by TEM (Scale bars = 1 μm).

**Supplementary Fig 2.**
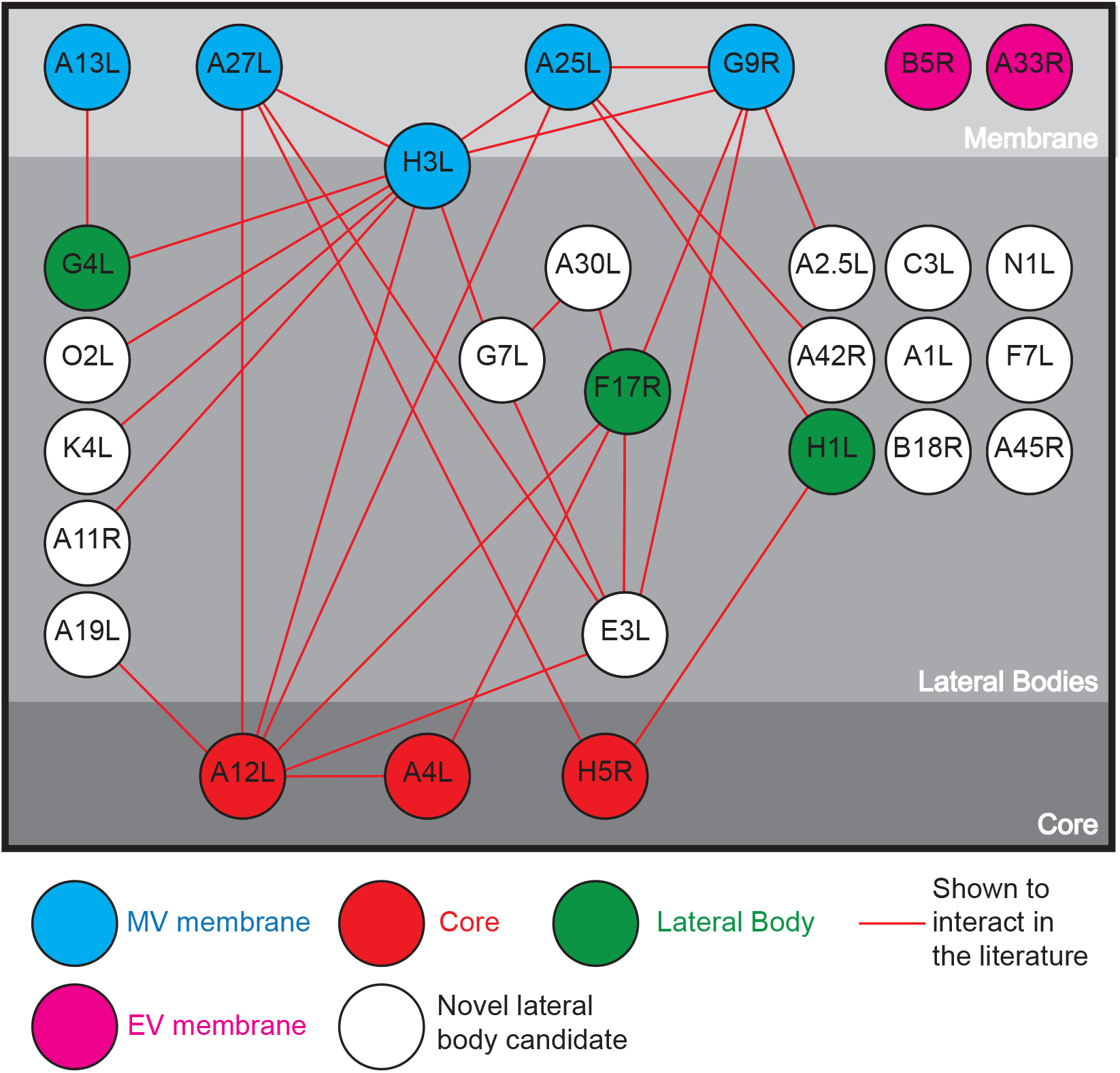
Schematic of VACV Membrane, LB and Core interactions. The VACV-MV protein-protein interaction network established by Mirzakhanyan and Gershon (59) was used to build a schematic of the interactions of the LB candidate proteins identified by this study. Proteins are colour-coded according to subviral location as indicated.

**Supplementary Fig 3.**
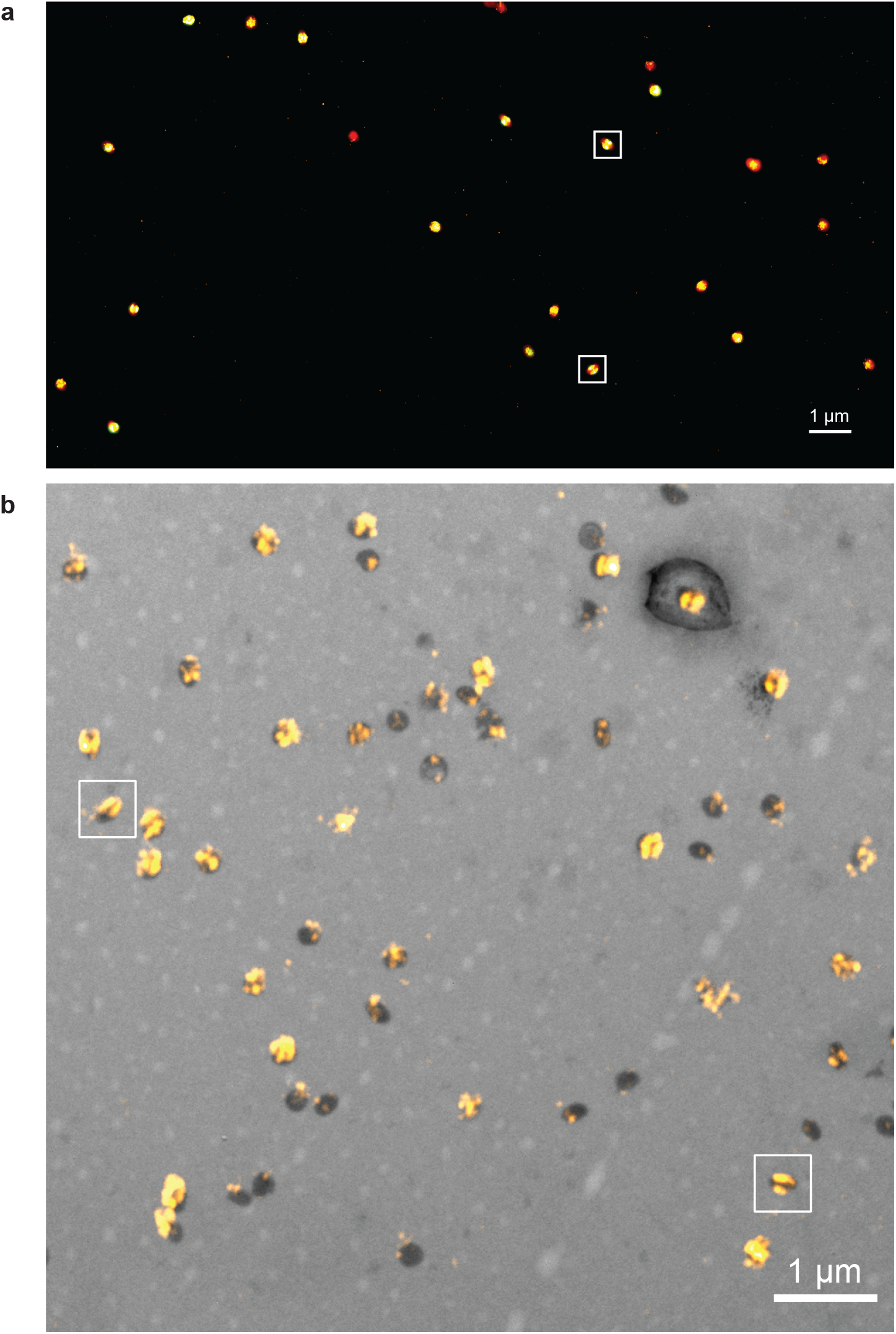
Overviews of Correlative SIM, STORM and EM. **(a)** Correlative SIM / STORM of WR mCherry-A4L A19L-EGFP virions. The two viral particles shown in Fig 3d are boxed. **(b)** Correlative STORM / EM of WR mCherry-A4L A19L-EGFP virions immunolabelled with anti-GFP nanobody. STORM images of the lateral body protein were registered with EM micrographs. The two viral particles shown in Fig 3e are boxed. Scale bars = 1 μm.

**Supplementary Fig 4.**
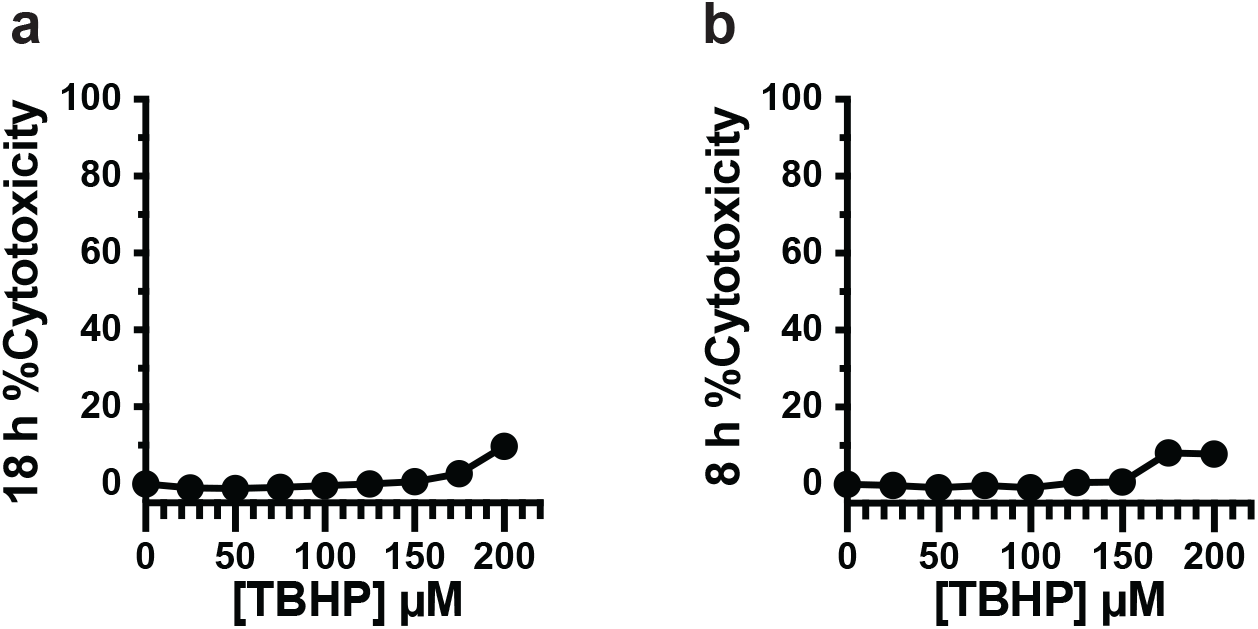
TBHP treatment is not cytotoxic. **(a)** LDH release assay for cytotoxicity of TBHP titration on A549s cells under the conditions used in Fig 4c. **(b)** LDH release assay for cytotoxicity of TBHP titration on A549s cells under the conditions used in Fig 4d & e. (n = 3 ± SEM).

## References

1. Mercer J, Schelhaas M, Helenius A. Virus entry by endocytosis. Annu Rev Biochem. 2010;79:803–33.

2. Marsh M, Helenius A. Virus entry: open sesame. Cell. 2006;124(4):729–40.

3. Tam JC, Jacques DA. Intracellular immunity: finding the enemy within--how cells recognize and respond to intracellular pathogens. J Leukoc Biol. 2014;96(2):233–44.

4. Brubaker SW, Bonham KS, Zanoni I, Kagan JC. Innate immune pattern recognition: a cell biological perspective. Annu Rev Immunol. 2015;33:257–90.

5. Beachboard DC, Horner SM. Innate immune evasion strategies of DNA and RNA viruses. Curr Opin Microbiol. 2016;32:113–9.

6. Garcia-Sastre A. Ten Strategies of Interferon Evasion by Viruses. Cell Host Microbe. 2017;22(2):176–84.

7. Bidgood SR, Mercer J. Cloak and Dagger: Alternative Immune Evasion and Modulation Strategies of Poxviruses. Viruses. 2015;7(8):4800–25.

8. Schmidt FI, Bleck CK, Reh L, Novy K, Wollscheid B, Helenius A, et al. Vaccinia virus entry is followed by core activation and proteasome-mediated release of the immunomodulatory effector VH1 from lateral bodies. Cell reports. 2013;4(3):464–76.

9. Yang L, Wang M, Cheng A, Yang Q, Wu Y, Jia R, et al. Innate Immune Evasion of Alphaherpesvirus Tegument Proteins. Front Immunol. 2019;10:2196.

10. Knipe S, Howley P. Fields Virology. Philadelphia, PA, USA.: Lippincott Williams & Wilkins; 2013.

11. Loret S, Guay G, Lippe R. Comprehensive characterization of extracellular herpes simplex virus type 1 virions. J Virol. 2008;82(17):8605–18.

12. Granzow H, Klupp BG, Mettenleiter TC. Entry of pseudorabies virus: an immunogold-labeling study. J Virol. 2005;79(5):3200–5.

13. Luxton GW, Haverlock S, Coller KE, Antinone SE, Pincetic A, Smith GA. Targeting of herpesvirus capsid transport in axons is coupled to association with specific sets of tegument proteins. Proc Natl Acad Sci U S A. 2005;102(16):5832–7.

14. Maurer UE, Sodeik B, Grunewald K. Native 3D intermediates of membrane fusion in herpes simplex virus 1 entry. Proc Natl Acad Sci U S A. 2008;105(30):10559–64.

15. Sodeik B, Ebersold MW, Helenius A. Microtubule-mediated transport of incoming herpes simplex virus 1 capsids to the nucleus. J Cell Biol. 1997;136(5):1007–21.

16. Dales S. The uptake and development of vaccinia virus in strain L cells followed with labeled viral deoxyribonucleic acid. J Cell Biol. 1963;18:51–72.

17. Ichihashi Y, Oie M, Tsuruhara T. Location of DNA-binding proteins and disulfide-linked proteins in vaccinia virus structural elements. J Virol. 1984;50(3):929–38.

18. Easterbrook KB. Controlled degradation of vaccinia virions in vitro: an electron microscopic study. J Ultrastruct Res. 1966;14(5):484–96.

19. Condit RC, Moussatche N, Traktman P. In a nutshell: structure and assembly of the vaccinia virion. Adv Virus Res. 2006;66:31–124.

20. Mercer J, Helenius A. Vaccinia virus uses macropinocytosis and apoptotic mimicry to enter host cells. Science. 2008;320(5875):531–5.

21. Schmidt FI, Bleck CK, Helenius A, Mercer J. Vaccinia extracellular virions enter cells by macropinocytosis and acid-activated membrane rupture. EMBO J. 2011;30(17):3647–61.

22. Huang CY, Lu TY, Bair CH, Chang YS, Jwo JK, Chang W. A novel cellular protein, VPEF, facilitates vaccinia virus penetration into HeLa cells through fluid phase endocytosis. J Virol. 2008;82(16):7988–99.

23. Yang Z, Moss B. Interaction of the vaccinia virus RNA polymerase-associated 94-kilodalton protein with the early transcription factor. J Virol. 2009;83(23):12018–26.

24. Laliberte JP, Weisberg AS, Moss B. The membrane fusion step of vaccinia virus entry is cooperatively mediated by multiple viral proteins and host cell components. PLoS Pathog. 2011;7(12):e1002446.

25. Locker JK, Griffiths G. An unconventional role for cytoplasmic disulfide bonds in vaccinia virus proteins. J Cell Biol. 1999;144(2):267–79.

26. Meade N, King M, Munger J, Walsh D. mTOR Dysregulation by Vaccinia Virus F17 Controls Multiple Processes with Varying Roles in Infection. J Virol. 2019;93(15).

27. Wickramasekera NT, Traktman P. Structure/Function analysis of the vaccinia virus F18 phosphoprotein, an abundant core component required for virion maturation and infectivity. J Virol. 2010;84(13):6846–60.

28. Mann BA, Huang JH, Li P, Chang HC, Slee RB, O’Sullivan A, et al. Vaccinia virus blocks Stat1-dependent and Stat1-independent gene expression induced by type I and type II interferons. J Interferon Cytokine Res. 2008;28(6):367–80.

29. Najarro P, Traktman P, Lewis JA. Vaccinia virus blocks gamma interferon signal transduction: viral VH1 phosphatase reverses Stat1 activation. J Virol. 2001;75(7):3185–96.

30. White CL, Senkevich TG, Moss B. Vaccinia virus G4L glutaredoxin is an essential intermediate of a cytoplasmic disulfide bond pathway required for virion assembly. J Virol. 2002;76(2):467–72.

31. White CL, Weisberg AS, Moss B. A glutaredoxin, encoded by the G4L gene of vaccinia virus, is essential for virion morphogenesis. J Virol. 2000;74(19):9175–83.

32. Senkevich TG, White CL, Koonin EV, Moss B. Complete pathway for protein disulfide bond formation encoded by poxviruses. Proc Natl Acad Sci U S A. 2002;99(10):6667–72.

33. Wink DA, Hines HB, Cheng RY, Switzer CH, Flores-Santana W, Vitek MP, et al. Nitric oxide and redox mechanisms in the immune response. J Leukoc Biol. 2011;89(6):873–91.

34. Khomich OA, Kochetkov SN, Bartosch B, Ivanov AV. Redox Biology of Respiratory Viral Infections. Viruses. 2018;10(8).

35. Checconi P, De Angelis M, Marcocci ME, Fraternale A, Magnani M, Palamara AT, et al. Redox-Modulating Agents in the Treatment of Viral Infections. Int J Mol Sci. 2020;21(11).

36. Paiva CN, Bozza MT. Are reactive oxygen species always detrimental to pathogens? Antioxid Redox Signal. 2014;20(6):1000–37.

37. Riva DA, de Molina MC, Rocchetta I, Gerhardt E, Coulombie FC, Mersich SE. Oxidative stress in vero cells infected with vesicular stomatitis virus. Intervirology. 2006;49(5):294–8.

38. Huang TT, Carlson EJ, Epstein LB, Epstein CJ. The role of superoxide anions in the establishment of an interferon-alpha-mediated antiviral state. Free Radic Res Commun. 1992;17(1):59–72.

39. Roederer M, Staal FJ, Raju PA, Ela SW, Herzenberg LA, Herzenberg LA. Cytokine-stimulated human immunodeficiency virus replication is inhibited by N-acetyl-L-cysteine. Proc Natl Acad Sci U S A. 1990;87(12):4884–8.

40. Staal FJ, Roederer M, Herzenberg LA, Herzenberg LA. Intracellular thiols regulate activation of nuclear factor kappa B and transcription of human immunodeficiency virus. Proc Natl Acad Sci U S A. 1990;87(24):9943–7.

41. Geiler J, Michaelis M, Naczk P, Leutz A, Langer K, Doerr HW, et al. N-acetyl-L-cysteine (NAC) inhibits virus replication and expression of pro-inflammatory molecules in A549 cells infected with highly pathogenic H5N1 influenza A virus. Biochem Pharmacol. 2010;79(3):413–20.

42. Mata M, Morcillo E, Gimeno C, Cortijo J. N-acetyl-L-cysteine (NAC) inhibit mucin synthesis and pro-inflammatory mediators in alveolar type II epithelial cells infected with influenza virus A and B and with respiratory syncytial virus (RSV). Biochem Pharmacol. 2011;82(5):548–55.

43. Vlahos R, Stambas J, Bozinovski S, Broughton BR, Drummond GR, Selemidis S. Inhibition of Nox2 oxidase activity ameliorates influenza A virus-induced lung inflammation. PLoS Pathog. 2011;7(2):e1001271.

44. Castro SM, Guerrero-Plata A, Suarez-Real G, Adegboyega PA, Colasurdo GN, Khan AM, et al. Antioxidant treatment ameliorates respiratory syncytial virus-induced disease and lung inflammation. Am J Respir Crit Care Med. 2006;174(12):1361–9.

45. Cho HY, Imani F, Miller-DeGraff L, Walters D, Melendi GA, Yamamoto M, et al. Antiviral activity of Nrf2 in a murine model of respiratory syncytial virus disease. Am J Respir Crit Care Med. 2009;179(2):138–50.

46. Speir E, Shibutani T, Yu ZX, Ferrans V, Epstein SE. Role of reactive oxygen intermediates in cytomegalovirus gene expression and in the response of human smooth muscle cells to viral infection. Circ Res. 1996;79(6):1143–52.

47. Tilton C, Clippinger AJ, Maguire T, Alwine JC. Human cytomegalovirus induces multiple means to combat reactive oxygen species. J Virol. 2011;85(23):12585–93.

48. Karupiah G, Xie QW, Buller RM, Nathan C, Duarte C, MacMicking JD. Inhibition of viral replication by interferon-gamma-induced nitric oxide synthase. Science. 1993;261(5127):1445–8.

49. Karupiah G, Chen JH, Nathan CF, Mahalingam S, MacMicking JD. Identification of nitric oxide synthase 2 as an innate resistance locus against ectromelia virus infection. J Virol. 1998;72(9):7703–6.

50. Ahn BY, Moss B. Glutaredoxin homolog encoded by vaccinia virus is a virion-associated enzyme with thioltransferase and dehydroascorbate reductase activities. Proc Natl Acad Sci U S A. 1992;89(15):7060–4.

51. Almazan F, Tscharke DC, Smith GL. The vaccinia virus superoxide dismutase-like protein (A45R) is a virion component that is nonessential for virus replication. J Virol. 2001;75(15):7018–29.

52. Senkevich TG, White CL, Weisberg A, Granek JA, Wolffe EJ, Koonin EV, et al. Expression of the vaccinia virus A2.5L redox protein is required for virion morphogenesis. Virology. 2002;300(2):296–303.

53. Satheshkumar PS, Olano LR, Hammer CH, Zhao M, Moss B. Interactions of the vaccinia virus A19 protein. J Virol. 2013;87(19):10710–20.

54. Satheshkumar PS, Weisberg AS, Moss B. Vaccinia virus A19 protein participates in the transformation of spherical immature particles to barrel-shaped infectious virions. J Virol. 2013;87(19):10700–9.

55. Rajagopal I, Ahn BY, Moss B, Mathews CK. Roles of vaccinia virus ribonucleotide reductase and glutaredoxin in DNA precursor biosynthesis. J Biol Chem. 1995;270(46):27415–8.

56. Guo T, Kouvonen P, Koh CC, Gillet LC, Wolski WE, Rost HL, et al. Rapid mass spectrometric conversion of tissue biopsy samples into permanent quantitative digital proteome maps. Nat Med. 2015;21(4):407–13.

57. Saveliev S, Bratz M, Zubarev R, Szapacs M, Budamgunta H, Urh M. Trypsin/Lys-C protease mix for enhanced protein mass spectrometry analysis. Nat Methods. 2013;10:i–ii.

58. Liu K, Lemon B, Traktman P. The dual-specificity phosphatase encoded by vaccinia virus, VH1, is essential for viral transcription in vivo and in vitro. J Virol. 1995;69(12):7823–34.

59. Mirzakhanyan Y, Gershon P. The Vaccinia virion: Filling the gap between atomic and ultrastructure. PLoS Pathog. 2019;15(1):e1007508.

60. Chung CS, Chen CH, Ho MY, Huang CY, Liao CL, Chang W. Vaccinia virus proteome: identification of proteins in vaccinia virus intracellular mature virion particles. J Virol. 2006;80(5):2127–40.

61. Resch W, Hixson KK, Moore RJ, Lipton MS, Moss B. Protein composition of the vaccinia virus mature virion. Virology. 2007;358(1):233–47.

62. Manes NP, Estep RD, Mottaz HM, Moore RJ, Clauss TR, Monroe ME, et al. Comparative proteomics of human monkeypox and vaccinia intracellular mature and extracellular enveloped virions. J Proteome Res. 2008;7(3):960–8.

63. Szklarczyk D, Gable AL, Lyon D, Junge A, Wyder S, Huerta-Cepas J, et al. STRING v11: protein-protein association networks with increased coverage, supporting functional discovery in genome-wide experimental datasets. Nucleic Acids Res. 2019;47(D1):D607–D13.

64. Poole LB. The basics of thiols and cysteines in redox biology and chemistry. Free Radic Biol Med. 2015;80:148–57.

65. Cao JX, Teoh ML, Moon M, McFadden G, Evans DH. Leporipoxvirus Cu-Zn superoxide dismutase homologs inhibit cellular superoxide dismutase, but are not essential for virus replication or virulence. Virology. 2002;296(1):125–35.

66. Teoh ML, Walasek PJ, Evans DH. Leporipoxvirus Cu,Zn-superoxide dismutase (SOD) homologs are catalytically inert decoy proteins that bind copper chaperone for SOD. J Biol Chem. 2003;278(35):33175–84.

67. Johnson GP, Goebel SJ, Perkus ME, Davis SW, Winslow JP, Paoletti E. Vaccinia virus encodes a protein with similarity to glutaredoxins. Virology. 1991;181(1):378–81.

68. Gray RD, Beerli C, Pereira PM, Scherer KM, Samolej J, Bleck CK, et al. VirusMapper: open-source nanoscale mapping of viral architecture through super-resolution microscopy. Sci Rep. 2016;6:29132.

69. Gray RDM, Mercer J, Henriques R. Open-source Single-particle Analysis for Super-resolution Microscopy with VirusMapper. J Vis Exp. 2017(122).

70. Culley S, Albrecht D, Jacobs C, Pereira PM, Leterrier C, Mercer J, et al. Quantitative mapping and minimization of super-resolution optical imaging artifacts. Nat Methods. 2018;15(4):263–6.

71. Gray RDM, Albrecht D, Beerli C, Huttunen M, Cohen GH, White IJ, et al. Nanoscale polarization of the entry fusion complex of vaccinia virus drives efficient fusion. Nat Microbiol. 2019;4(10):1636–44.

72. Gray R, Albrecht D. Super-resolution Microscopy of Vaccinia Virus Particles. Methods Mol Biol. 2019;2023:255–68.

73. Huttunen M, Mercer J. Quantitative PCR-Based Assessment of Vaccinia Virus RNA and DNA in Infected Cells. Methods Mol Biol. 2019;2023:189–208.

74. Ichihashi Y. Extracellular enveloped vaccinia virus escapes neutralization. Virology. 1996;217(2):478–85.

75. Smiley JR. Herpes simplex virus virion host shutoff protein: immune evasion mediated by a viral RNase? J Virol. 2004;78(3):1063–8.

76. Risco C, Rodriguez JR, Lopez-Iglesias C, Carrascosa JL, Esteban M, Rodriguez D. Endoplasmic reticulum-Golgi intermediate compartment membranes and vimentin filaments participate in vaccinia virus assembly. J Virol. 2002;76(4):1839–55.

77. Yoder JD, Chen TS, Gagnier CR, Vemulapalli S, Maier CS, Hruby DE. Pox proteomics: mass spectrometry analysis and identification of Vaccinia virion proteins. Virol J. 2006;3:10.

78. Wood JJ, White IJ, Mercer J. Acrylamide Inhibits Vaccinia Virus Through Vimentin-independent Anti-Viral Granule Formation. bioRXiv. 2020; https://doi.org/10.1101/2020.07.30.228858.

79. Takahashi T, Oie M, Ichihashi Y. N-terminal amino acid sequences of vaccinia virus structural proteins. Virology. 1994;202(2):844–52.

80. Szajner P, Jaffe H, Weisberg AS, Moss B. A complex of seven vaccinia virus proteins conserved in all chordopoxviruses is required for the association of membranes and viroplasm to form immature virions. Virology. 2004;330(2):447–59.

81. Condit RC, Moussatche N. The vaccinia virus E6 protein influences virion protein localization during virus assembly. Virology. 2015;482:147–56.

82. Moussatche N, Condit RC. Fine structure of the vaccinia virion determined by controlled degradation and immunolocalization. Virology. 2015;475:204–18.

83. Mercer J, Traktman P. Genetic and cell biological characterization of the vaccinia virus A30 and G7 phosphoproteins. J Virol. 2005;79(11):7146–61.

84. Novy K, Kilcher S, Omasits U, Bleck CKE, Beerli C, Vowinckel J, et al. Proteotype profiling unmasks a viral signalling network essential for poxvirus assembly and transcriptional competence. Nat Microbiol. 2018;3(5):588–99.

85. Dobson BM, Tscharke DC. Redundancy complicates the definition of essential genes for vaccinia virus. J Gen Virol. 2015;96(11):3326–37.

86. Sumner RP, Maluquer de Motes C, Veyer DL, Smith GL. Vaccinia virus inhibits NF-kappaB-dependent gene expression downstream of p65 translocation. J Virol. 2014;88(6):3092–102.

87. Rizopoulos Z, Balistreri G, Kilcher S, Martin CK, Syedbasha M, Helenius A, et al. Vaccinia Virus Infection Requires Maturation of Macropinosomes. Traffic. 2015.

88. Yakimovich A, Huttunen M, Zehnder B, Coulter LJ, Gould V, Schneider C, et al. Inhibition of Poxvirus Gene Expression and Genome Replication by Bisbenzimide Derivatives. J Virol. 2017;91(18).

89. Agrotis A, Pengo N, Burden JJ, Ketteler R. Redundancy of human ATG4 protease isoforms in autophagy and LC3/GABARAP processing revealed in cells. Autophagy. 2019;15(6):976–97.

90. Eng JK, Jahan TA, Hoopmann MR. Comet: an open-source MS/MS sequence database search tool. Proteomics. 2013;13(1):22–4.

91. Keller A, Nesvizhskii AI, Kolker E, Aebersold R. Empirical statistical model to estimate the accuracy of peptide identifications made by MS/MS and database search. Anal Chem. 2002;74(20):5383–92.

92. Choi M, Chang CY, Clough T, Broudy D, Killeen T, MacLean B, et al. MSstats: an R package for statistical analysis of quantitative mass spectrometry-based proteomic experiments. Bioinformatics. 2014;30(17):2524–6.

93. Benjamini Y, Hochberg Y. Controlling the False Discovery Rate - a Practical and Powerful Approach to Multiple Testing. J R Stat Soc B. 1995;57(1):289–300.

94. Chakrabarti S, Sisler JR, Moss B. Compact, synthetic, vaccinia virus early/late promoter for protein expression. Biotechniques. 1997;23(6):1094–7.

95. Giotis ES, Laidlaw SM, Bidgood SR, Albrecht D, Burden JJ, Robey RC, et al. Modulation of early host innate immune response by a Fowlpox virus (FWPV) lateral body protein. bioRxiv. 2020(https://doi.org/10.1101/2020.10.02.324418).

